# Male mice with large inversions or deletions of X-palindrome arms are fertile and express their associated genes post-meiosis

**DOI:** 10.1101/245860

**Authors:** Alyssa N. Kruger, Quinn Ellison, Michele A. Brogley, Emma R. Gerlinger, Jacob L. Mueller

## Abstract

Large (>10 kb) palindromic sequences are enriched on mammalian sex chromosomes. In mice, these palindromes harbor gene families (≥2 gene copies) expressed exclusively in post-meiotic testicular germ cells, at a time when most single-copy sex-linked genes are transcriptionally repressed. This distinct expression pattern led to the hypothesis that containment within palindrome structures or having ≥2 gene enables post-meiotic gene expression. We tested these two hypotheses by using CRISPR to precisely engineer large (10’s of kb) inversions and deletions of X chromosome palindrome arms for two regions carrying the mouse *4930567H17Rik* and *Mageb5* gene families. We found that *4930567H17Rik* and *Mageb5* gene expression is unaffected in mice carrying palindrome arm inversions, suggesting that palindromic structure is not important for mediating palindrome-associated gene expression. We also found that *4930567H17Rik* and *Mageb5* gene expression is reduced by half in mice carrying palindrome arm deletions, allowing us to test whether palindrome-associated genes are sensitive to reduced expression levels resulting in spermatogenic defects. Male mice carrying palindrome arm deletions of *4930567H17Rik* or *Mageb5*, however, are fertile, have normal testis histology, and show no aberrations in spermatogenic cell population frequencies via FACS quantification. Together, these findings suggest that large palindromic structures on the sex chromosomes are not necessary for their associated genes to evade post-meiotic transcriptional repression and that these genes are not sensitive to reduced expression levels. Large sex chromosome palindromes may thus be important for other reasons, such as the long-term evolutionary stability of their associated gene families.

## Introduction

In humans and mice, the sex chromosomes are enriched for large (>10kb), nearly identical (>99% nucleotide identity) segmental duplications in palindromic orientation^1-4^. In mice, genes harbored within large X chromosome palindromes are expressed predominantly or exclusively in post-meiotic testicular germ cells^3^. This specific expression pattern is surprising, because most single-copy X-linked genes are transcriptionally silenced post-meiosis^5-8^. The mechanism by which palindrome-associated genes escape transcriptional repression is unknown; however, two hypotheses have been suggested to explain this distinct expression pattern. First, palindromes may form secondary structures (e.g. palindrome arms pairing to form a hairpin) enabling their associated genes to evade transcriptional repression^3^. Intrachromosomal synapsis of palindrome arm pairing could facilitate the evasion of post-meiotic gene repression, which itself is a consequence of asynapsis-triggered meiotic sex chromosome inactivation^9-11^. Second, X-palindromic genes may be sensitive to reduced expression levels and thus require ≥2 gene copies^3^. Consistent with this, the mouse X chromosome carries other non-palindromic genes that are in multiple copies and are expressed specifically in post-meiotic cells^3^. To test the two hypotheses, individual palindrome arms must be genetically manipulated, in vivo.

To rigorously test whether palindrome structure or gene copy number is important for post-meiotic expression, we genetically dissected two mouse X-palindromes. We utilized CRISPR to generate large-scale (10’s of kb) inversions and deletions in mice of two X-palindrome arms harboring the *4930567H17Rik* and *Mageb5* (*Melanoma antigen gene family member b5*) gene families. We chose these two X-palindromes because of their canonical features of palindromes across mammals; they have >99% percent nucleotide identity between the two arms, the arms are >10kb, and they harbor a gene family expressed specifically in post-meiotic testicular germ cells. We also selected these two gene families because they have sequence family variants between the two gene copies, enabling detection of palindrome arm-specific expression. We found that for the *4930567H17Rik* and *Mageb5* palindromic gene families, palindrome structure is not necessary for regulating their associated post-meiotic gene expression. We observed that deletion of a single palindrome arm, for both the *4930567H17Rik* and *Mageb5* gene families, reduces gene expression levels by half; however, reduced expression levels did not lead to male infertility or spermatogenic defects in either case. This suggests that palindromes enrichment on the sex chromosomes is important for other reasons and that there are alternative, unknown mechanisms for palindrome-associated genes to evade post-meiotic repression.

## Materials and Methods

### Generation of 4930567H17Rik and Mageb5 palindrome arm inversions and deletions

To generate mice carrying palindrome arm inversions and deletions we used CRISPR with dual sgRNAs (single guide RNAs). sgRNAs were designed to unique sequences flanking the targeted palindrome arm, as close to the edge of the arm as possible. Since sgRNA cutting efficiency varies between sgRNAs, we tested their activity via pronuclear injection of an sgRNA/Cas9 expression plasmid, pSpCas9(BB)-2A-GFP (pX458)^12^ in mouse zygotes. The surviving mouse zygotes were allowed to develop into 64-128 cell blastocysts and the cutting efficiency of each sgRNA was determined by PCR of the cut sties and Sanger sequencing of purified PCR products to identify local edits at the target sites.

Once active sgRNAs were identified for both sides of the targeted palindrome arm, two pX458M plasmids encoding the sgRNAs and Cas9 together with a single-stranded oligonucleotide were pronuclear injected into hybrid C57BL/6JxSJL hybrid mouse zygotes. We added single-stranded oligonucleotides with sequence homology flanking the junction boundaries, to promote either inversions or deletions. The zygotes were generated from mating C57BL/6J females to C57BL/6JxSJL males so that all targeted X chromosomes were C57BL/6J. Blastocysts from the pronuclear injection where implanted into pseudopregnant females. Genomic DNA from resulting pups was screened via PCR and Sanger sequencing for the inversion and deletion junctions. At least two independent mouse lines were obtained for inversions and deletions of the *4930567H17Rik* and *Mageb5* palindrome arms. Male and female mice were able to germline transmit both the *4930567H17Rik* and *Mageb5* deletions and inversions, thus their overall health was unaffected by our CRISPR-mediated chromosome engineering.

### Mice and testis sample collection

All mice carrying an inversion or deletion of a single palindrome arm were backcrossed to C57BL/6J (N2-N7). Backcrossing was performed to minimize any possible CRISPR-mediated off-target effects. For assessing fecundity (average litter size per male), we mated mutant males to CD1 females. Testes were collected from 2-6 month old males for all experiments. We used wild-type littermate controls whenever possible and if not available, we used age-matched controls from the same breeding line. The alleles for the first mouse lines of each type were named *4930567H17Rik*^*InvArm1*^, *4930567H17Rik*^*DelArm1*^, *Mageb5*^*InvArm1*^, *Mageb5*^*DelArm1*^ with respective registered symbols Del(X4930567H17Rik)1Jbmu, In(X4930567H17Rik)1Jbmu, Del(XMageb5)1Jbmu, In(XMageb5)1Jbmu. All experiments performed on mice were approved by the University of Michigan Committee on Use and Care of Animals, and all experiments followed the National Institutes of Health Guidelines of the Care and Use of Experimental Animals.

### Preparation of adult testis cDNA for qPCR

Intron-spanning primers, when possible, were used to perform qPCR on adult testis cDNA preparations. One testis per mouse was used to isolate RNA using Trizol (Invitrogen) following the manufacturers recommendations. 5 μg of total RNA was DNase treated with TurboDNase (Ambion) and reversed transcribed using Superscript II (Invitrogen) and oligonucleotide (dT) primers following the manufacturers protocol. qPCR was performed in triplicate using Power SYBR Green master mix (Thermo Fisher Scientific) on a 7500 Real-time PCR thermalcycler (Applied Biosystems). *Trim42* (*Tripartite motif-containing 42*) was used as a normalization control, because it is expressed specifically in the same post-meiotic testicular cells and at similar levels as *4930567H17Rik* and *Mageb5*. The Delta-delta Ct method, with *Trim42* as the normalization control was used determine gene expression differences.

### Testis Histology

Testes were fixed overnight in Bouin’s solution, paraffin embedded, sectioned to 5 μm, and stained with Periodic acid Schiff (PAS) and hematoxylin. Sections were visualized under a light microscope and specific germ cell populations were identified by their location within a tubule, nuclear size, and chromatin pattern^13^.

### FACs-based estimates of round spermatid frequencies

We largely followed a previously published protocol^14^ to isolate round spermatids (1n) and spermatocytes (4n). Briefly, we disassociated cells from a single testis by enzymatic treatment with Collagenase type I, DNase I (Worthington Biochemical Corporation), and Trypsin (Life Technologies). The cell suspension was passed through cell strainers (100 μm and 40 μm) and incubated with Hoechst 33342 (Life Technologies) for DNA content and propidium iodide (Acros Organics) for cell viability. Cell sorting was performed on a FACSAria II cell sorter (BD Biosciences). The purity of each sort was determined via fluorescence microscopy visual inspection of 100 cells morphology and nuclear staining with DAPI.

### RNA-seq analyses

RNA-seq analyses were conducted by analyzing previously published datasets. Specifically, mouse tissue panel data was analyzed from GSE41637^15^, ovary data from GSE43520^16^ and sorted testicular germ cell populations from GSE49624^17^. Alignments were performed using Tophat^18^ with the mm10 mouse reference genome, a refFlat file with RefSeq gene annotations and --max-multihits set to 240; otherwise standard default parameters were used. We used Cufflinks^18^ using the refFlat gene annotation file to estimate expression levels as fragments per kilobase per millions of mapped fragments (FPKM).

### Dot plots

Self-symmetry triangular dot plots that show repeats within a sequenced region were generated from a custom Perl script that can be found at http://pagelab.wi.mit.edu/material-request.html.

## Results

### The mouse X chromosome harbors eight singleton palindromes

Large palindromes on mammalian sex chromosomes are typically found as isolated pairs of palindrome arms (singleton palindromes) or in complex arrays of palindromes. We investigated singleton palindromes, because they are more commonly found across mammalian sex chromosomes and can be genetically manipulated in vivo more precisely. Of the eight singleton palindromes on the mouse X chromosome (Table 1 and Figure 1A), we selected two harboring the *4930567H17Rik* and *Mageb5,* because they share canonical features of sex chromosome palindromes: >10kb, >99% nucleotide identity between palindrome arms, harbor genes expressed specifically in testicular germ cells, and have a spacer sequence between the palindrome arms (Table 1 and Figure 1B). Additionally, the palindrome carrying the 4930567H17Rik gene family has the longest palindrome arm (65kb), for a singleton palindrome, which if we can delete and invert in mice, will serve as a proof of principle for the manipulation of shorter palindrome arms.

**Table 1.**
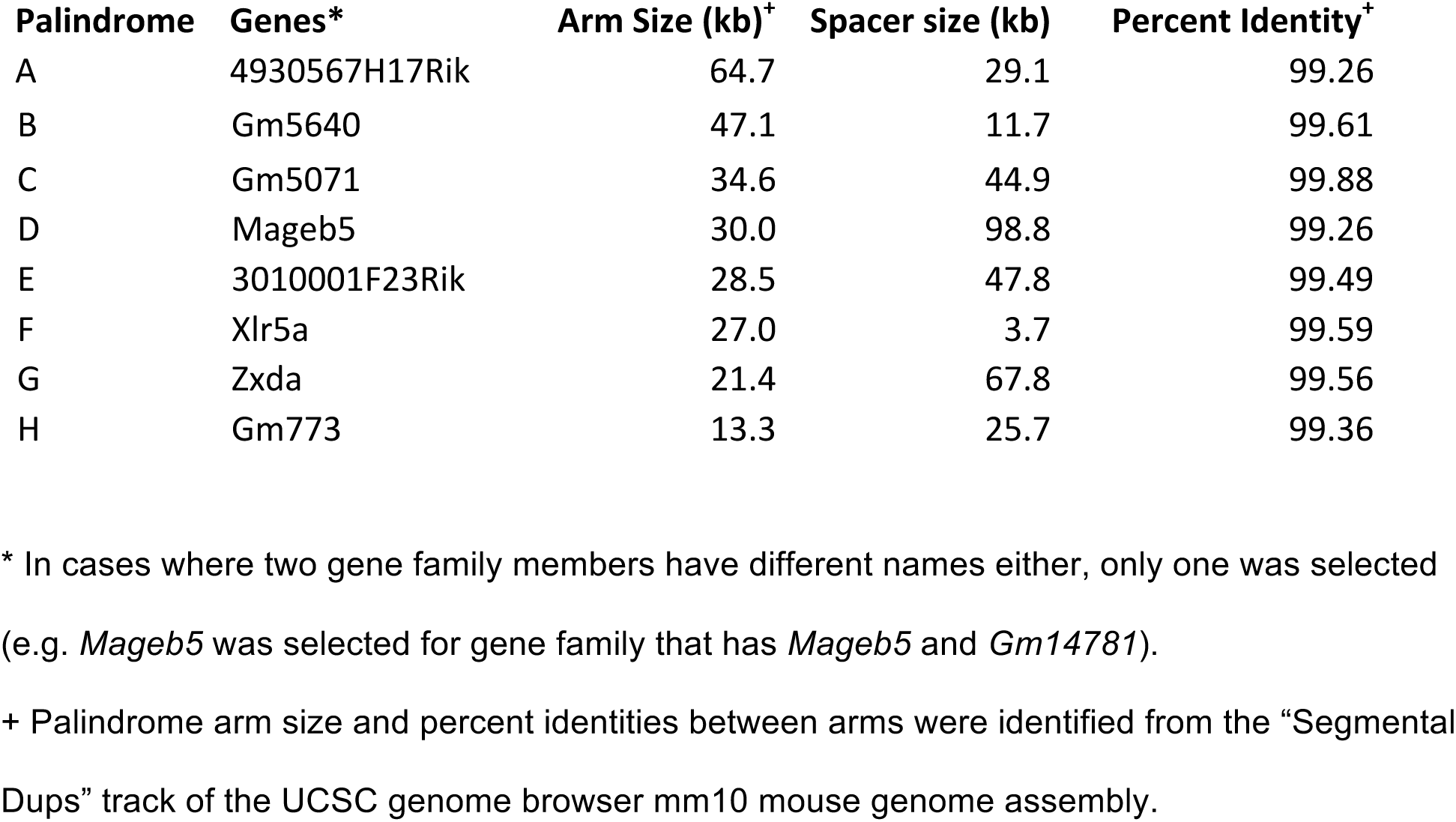

**Figure 1.**
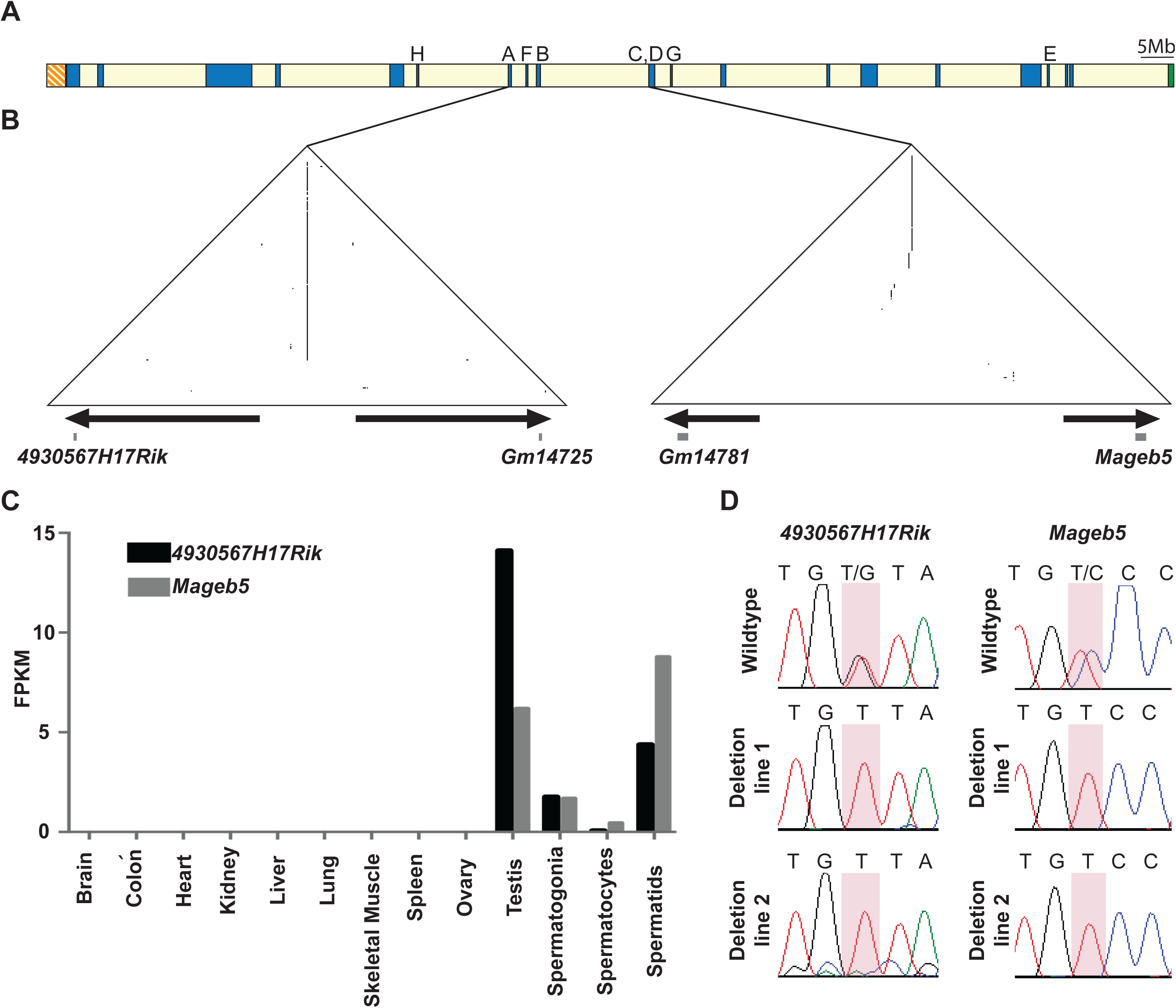
Singleton palindromes on the mouse X chromosome. **A**) The location of the eight singleton palindromic regions on the mouse X chromosome are labeled alphabetically, the sequence features of which can be found in Table 1. **B**) Self-symmetry triangular dot plots of the two singleton X-palindromes to be studied, carrying the *4930567H17Rik* and *Mageb5* gene families, respectively. Each dot plot represents the palindromic X chromosome sequence (*4930567H17Rik* = chrX:70385921-70553920 and *Mageb5*= chrX:91624421-91790420) plotted against itself with a sliding window of 100 nucleotides (step size=1 nucleotide). When the window of 100 nucleotides is identical to the sequence it is compared to, a dot is plotted. Segmental duplications in an inverted orientation are visualized as vertical lines. A visual representation of the palindrome arms (arrows) and the >99% identical genes harbored within them (squares) is plotted at the base of the triangular plots across the 168 kb *4930567H17Rik* and 166kb *Mageb5* palindromes. **C**) Expression levels of *4930567H17Rik* and *Mageb5* genes in adult tissues and sorted spermatogenic populations, shown as FPKMs (number of fragments per kilobase per million mapped fragments). **D**) Sanger sequencing of RT-PCR products displaying the sequence family variants that distinguish the two palindromic gene copies. In *4930567H17Rik*^*DelArm/Y*^and *Mageb5*^*DelArm/Y*^ mice, expression is detected only from the remaining copy.

We wanted to ensure that the *4930567H17Rik* and *Mageb5* gene families are expressed exclusively in post-meiotic round spermatids and that both copies are expressed. By reanalyzing previously published RNA-seq datasets, we find that both gene families are expressed exclusively in round spermatids (Figure 1C). We consider the low levels of expression in spermatogonia to be due to contamination of round spermatids in the spermatogonial populations during sorting. To determine if both *4930567H17Rik* and *Mageb5* gene copies are expressed, we utilized individual nucleotide differences between gene copies. Sequencing of RT-PCR products for both *4930567H17Rik* and *Mageb5* show that both gene copies are expressed (Figure 1D). Having confirmed that both gene copies are expressed exclusively in post-meiotic round spermatids, we proceeded to delete and invert individual palindrome arms to assess the importance of palindrome structure and gene copy number.

### Generation of mice carrying precise inversions and deletions of individual X-palindrome arms via CRISPR

We utilized CRISPR/Cas9 technology to precisely invert and delete large X-palindromes arms in mice. Pronuclear injection of mouse zygotes with single guide RNAs (sgRNAs), targeting unique flanking regions of each palindrome arm, and a single stranded oligonucleotide donor enabled us to generate large (29 kb and 65 kb) inversions and deletions of individual arms for the *4930567H17Rik* and *Mageb5* X-palindromes (Figure 2A). The single stranded oligonucleotide donors were used to promote inversions and deletions. We screened founder mouse lines with all combinations of primers flanking the sgRNA cut sites in order to detect and distinguish inversions and deletions (Figure 2B). We validated inversion and deletion junctions by Sanger sequencing (Figure 2C). After ~300 pronucelar injections we obtained 2 and 3 independent mouse lines carrying deletions of a single palindrome arm for the *4930567H17Rik* and *Mageb5* gene families, respectively. Similarly, after ~300 pronuclear injections we obtained 4 and 2 independent mouse lines carrying inversions of a single palindrome arm for the *4930567H17Rik* and *Mageb5* gene families, respectively. The independently obtained inversion and deletion junctions differed from each other by only a few nucleotides. We also confirmed the deletions via Sanger sequencing of RT-PCR products to show that only a single sequence family variant was expressed (Figure 1D) in each mouse line. We selected two deletion and two inversion independent mouse lines for each X-palindromic region. Throughout this study, the genotypes for the eight deletion and inversion carrying male lines are *4930567H17RiK*^*InvArm/Y*^, *4930567H17Rik*^*DelArm/Y*^, *Mageb5*^*InvArm/Y*^, and *Mageb5*^*DelArm/Y*^, followed by a 1 or 2 depending on the mouse line. Our dual sgRNAs combined with single stranded oligonucleotides approach was successful in generating multiple independent mouse lines carrying 29 kb and 65kb inversions and deletions of single palindrome arms from a single round of ~300 pronuclear injections.

**Figure 2.**
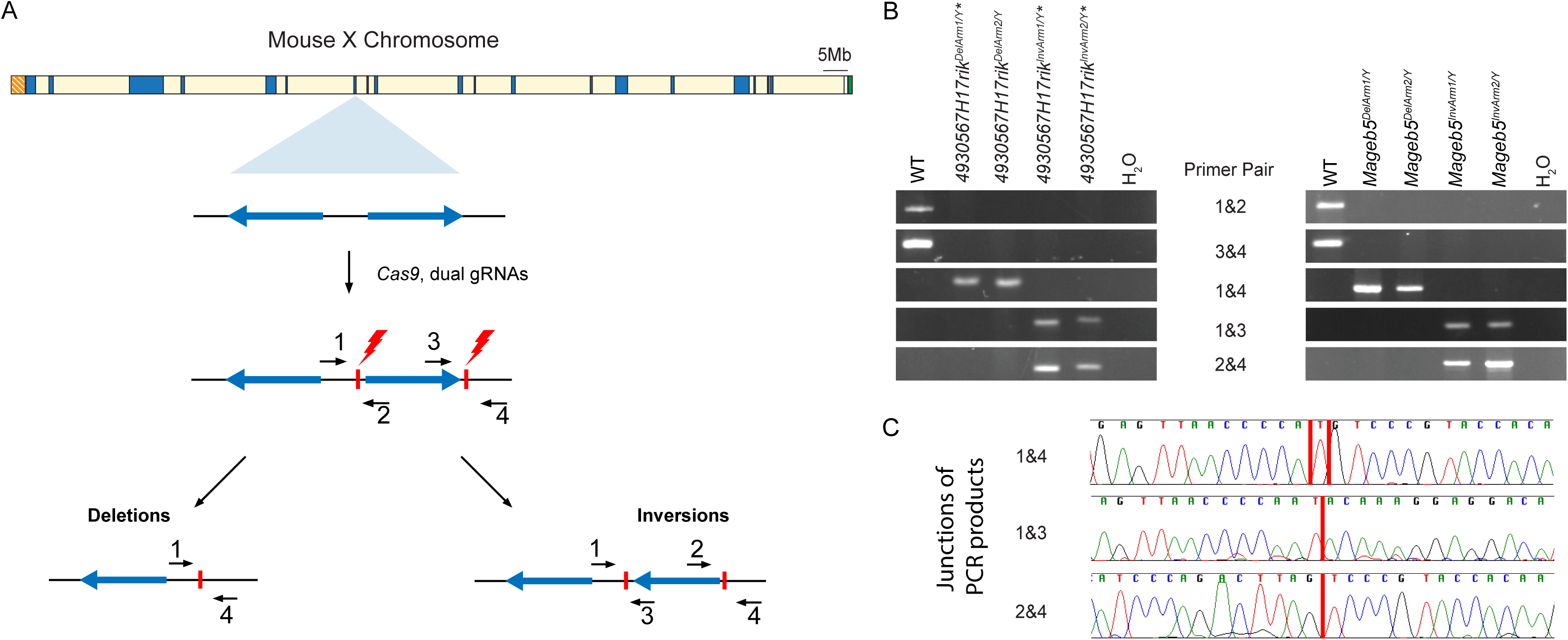
CRISPR strategy to generate large inversions and deletions of individual palindrome arms. **A)** Schematic of the mouse X chromosome with a diagram of a singleton palindrome shown below. Palindrome arms are shown as blue arrows. Single gRNA sites are shown as vertical red lines and primers to detect inversions and deletions are shown as black arrows. **B)** PCR genotyping of DNA from the two independent mouse lines for each of the deletions and inversions of *4930567H17Rik* and *Mageb5* palindrome arms. Numbered primers from panel A were used to amplify deletion (1+4) and inversion (1+3, 2+4) junctions. WT = wild-type **C)** Representative example of Sanger sequencing the deletion and inversion junctions for *4930567H17Rik* palindrome arm rearrangements. Junction sites are shown with a vertical red line. The mm10 coordinates for the sequence removed in the *4930567H17Rik*^*DelArm1/Y*^ line are ChrX:70389542-70457358 and coordinates for the sequence inverted in the *4930567H17RiK*^*InvArm1/Y*^ are ChrX:70389544-70457357.

### Disruption of palindrome structure, via inverting a single palindrome arm, does not affect the gene expression levels of the palindrome associated gene family

With our generation of mice carrying precise inversions of two different X-palindrome arms that disrupt palindrome structure, we tested whether palindrome structure is necessary for facilitating post-meiotic gene expression. If palindrome structure is necessary for post-meiotic gene expression, then we expected to abolish gene expression of *4930567H17Rik* and *Mageb5* in *4930567H17RiK*^*InvArm/Y*^ and *Mageb5*^*InvArm/Y*^ mice. Using gene-specific primers for *4930567H17Rik* and *Mageb5*, we compared their testis gene expression levels via quantitative RT-PCR (qRT-PCR) in *4930567H17RiK*^*InvArm/Y*^ and *Mageb5*^*InvArm/Y*^ mice carrying to wild-type controls and normalized to *Trim42*. After normalization, we find that *4930567H17RiK*^*InvArm/Y*^ and *Mageb5*^*InvArm/Y*^ mice, across two independent mouse lines, express their associated genes at similar levels to wild-type mice (Figure 3A). Consistent with their similar gene expression levels, *4930567H17RiK*^*InvArm/Y*^ and *Mageb5*^*InvArm/Y*^ are fertile and do no exhibit overt spermatogenic defects. This suggests that palindrome structure is not necessary for the expression of the *4930567H17Rik* and *Mageb5* X-palindrome genes.

**Figure 3.**
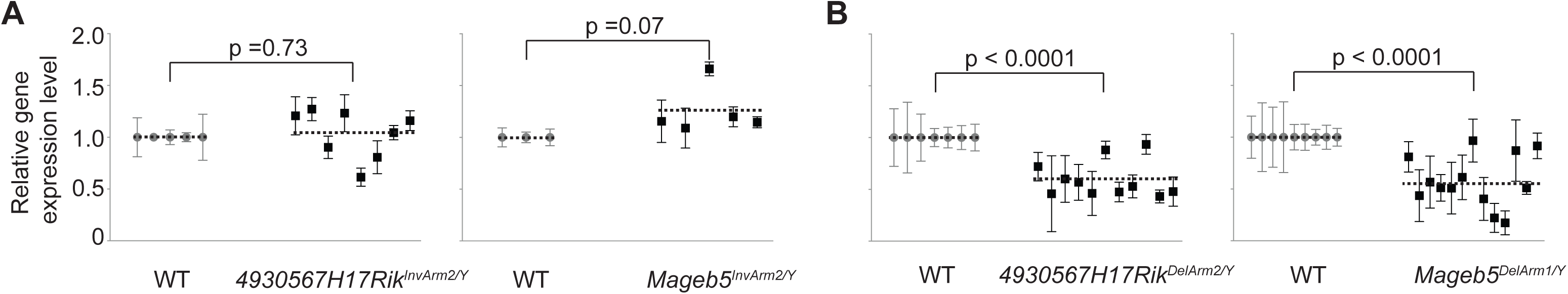
Male mice carrying *4930567H17Rik* and *Mageb5* palindrome arm inversions or deletions express their associated genes at wild-type and half of wild-type levels, respectively. **A)** *4930567H17Rik* and *Mageb5* gene expression levels from testes of *4930567H17Rik*^*InvArm/Y*^and *Mageb5*^*InvArm/Y*^, respectively. **B)** *4930567H17Rik* and *Mageb5* geneexpression levels from testes of *4930567H17Rik*^*DelArm/Y*^ and *Mageb5*^*DelArm/Y*^ mice, respectively. For **A** and **B**, the gene expression values for each individual mouse (each dot) are normalized to wild-type expression (WT= 1). All expression values are normalized to *Trim42*, which is expressed specifically in round spermatids. Error bars represent the standard deviation of technical replicates. Gene expression levels from independent mouse lines. Welch’s two-tailed t-tests were used to determine significance.

### Deleting a single palindrome arm reduces the gene expression level of the palindrome-associated gene family by half

Using mice carrying single palindrome arm deletions, we reduced the gene copy number of the palindrome associated *4930567H17Rik* and *Mageb5* X-linked genes by half (2 gene copies to 1). Concordantly, we expected *4930567H17Rik* and *Mageb5* gene expression levels would drop by half in *4930567H17Rik*^*DelArm/Y*^ and *Mageb5*^*DelArm/Y*^ mice. Using gene-specific primers for *4930567H17Rik* and *Mageb5*, we compared the testis gene expression levels of *4930567H17Rik*^*DelArm/Y*^ and *Mageb5*^*DelArm/Y*^ mice to wild-type controls and normalized to *Trim42*. After normalization, we indeed find that *4930567H17Rik*^*DelArm/Y*^ and *Mageb5*^*DelArm/Y*^ mice, across two independent mouse lines, express their associated genes at approximately half the levels of wild-type males (Figure 3B). The reduction of gene expression by half allows us to test whether the *4930567H17Rik* and *Mageb5* gene families are sensitive to reduced gene expression levels, resulting in males that are infertile with defects in post-meiotic spermatogenic development.

### Male mice carrying deletions of individual palindrome arms do not exhibit defects in fecundity, testis histology, or numbers of round spermatids

We performed a systematic characterization of fecundity and post-meiotic spermatogenic development in *4930567H17Rik*^*DelArm/Y*^ and *Mageb5*^*DelArm/Y*^ mice. We found that *4930567H17Rik*^*DelArm/Y*^ and *Mageb5*^*DelArm/Y*^ mice are fertile and produce litter sizes that are similar to wild-type controls (Figure 4A). To detect potential defects in post-meiotic spermatid development, we examined testis histological sections of *4930567H17Rik*^*DelArm/Y*^ and *Mageb5*^*DelArm/Y*^ mice. We did not observe defects in spermatid morphology, formation of the acrosome, initiation of spermatid elongation or their formation or the presence of abnormal elongated spermatids in the epithelium. To assess whether the number of round spermatids were affected in *4930567H17Rik*^*DelArm/Y*^ and *Mageb5*^*DelArm/Y*^ mice, we quantified the number of round spermatids per testis as the ratio of round spermatids/spermatocytes (control) via FACs. We find that the ratio of round spermatids/spermatoyctes of *4930567H17Rik*^*DelArm/Y*^ and *Mageb5*^*DelArm/Y*^ mice is similar to wild-type males (Figure 4B), which is consistent with their testis weights also being similar. Altogether, *4930567H17Rik*^*DelArm/Y*^ and *Mageb5*^*DelArm/Y*^ mice do not exhibit detectable defects in fecundity and post-meiotic spermatid development.

**Figure 4.**
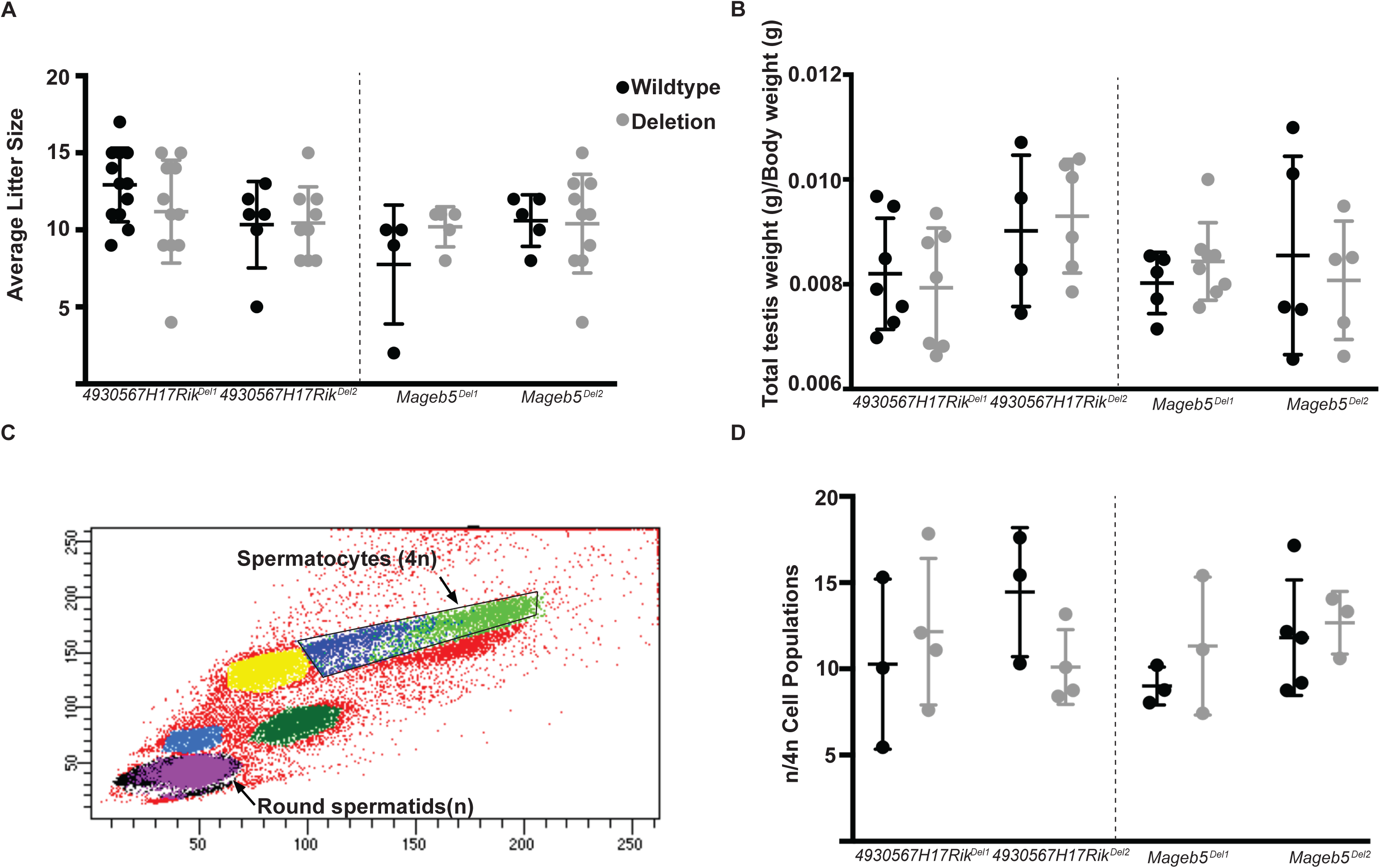
Male mice carrying *4930567H17Rik* and *Mageb5* palindrome arm deletions are fertile and do not display defects in spermatogenic cell population frequencies. **A)** Multiple males from each *4930567H17Rik*^*DelArm/Y*^ and *Mageb5*^*DelArm/Y*^ line and control males were mated to multiple CD1 females to assess fertility and fecundity. **B)** Total testis weight (g) from *4930567H17Rik*^*DelArm/Y*^and *Mageb5*^*DelArm/Y*^ lines were normalized to total body weight (g).Deletion males were compared to wild-type littermates. **C)** Representative FACS plot of spermatogenic cell populations with round spermatids (1n) and spermatocytes (4n) populations indicated by black arrows. **D)** The frequency of post-meiotic round spermatids was assessed by normalizing the percentage of cells in the post-meiotic round spermatid (1n) gate to the percentage of cells in the spermatocytes (4n).

## Discussion

Our finding that palindrome structure is not necessary for facilitating post-meiotic gene expression for *4930567H17Rik* and *Mageb5* gene families leaves open the question as to why palindromes are heavily enriched on both the mammalian X and Y chromosomes. It is possible that palindromes, both on the X and Y chromosomes, are necessary for long-term evolutionary stability of the genes they harbor in order to rapidly purge deleterious mutations via gene conversion^19^.

Our findings also show that the *4930567H17Rik* and *Mageb5* gene families are not sensitive to reduced gene expression levels, when a single palindrome arm deletion reduces the gene copy number from 2 to 1. This, together with our findings on palindrome structures, suggests there are alternative mechanisms for X-palindromic genes to be expressed on the otherwise transcriptionally repressed X chromosome. There are a small number of X-linked single-copy genes expressed in round spermatids^6,20^, indicating that multiple gene copies are not a strict requirement for post-meiotic gene expression from the sex chromosomes. Thus, specific enhancers and transcription factors may have evolved to overcome the transcriptional repression associated with the rest of the sex chromosomes. Two potential transcription factors that facilitate post-meiotic sex-linked gene expression are Heat Shock Transcription Factor 1 (HSF1), which localizes to sex chromatin^21^ and HSF2, which preferentially binds chromatin of Y-palindromic genes^22^. Consistent with this, the testis is known to have specialized transcription regulation strategies in post-meiotic testicular germ cells^23^ and appears to have evolved a unique mechanism for palindromic and multicopy X- and Y-linked genes.

## Acknowledgements

We thank Dirk de Rooij, Thomas Wilson and Michael Pihalja for technical advice, the University of Michigan Transgenic Animal Model Core for performing pronuclear injections to generate the *4930567H17Rik* and *Mageb5* palindrome arm deletion and inversion mouse lines, Flow Cytometry Core in order for us to perform FACs, DNA Sequencing Core for performing Sanger Sequencing, and Cancer Center Tissue Core for generating testis histological sections. These studies were supported by National Institutes of Health grants R00HD064753 to JLM, T32GM007544 to ANK and T32HD079342 to QE and a National Science Foundation Graduate Research Fellowship to ANK.

